# MultiDCoX: Multi-factor Analysis of Differential Coexpression

**DOI:** 10.1101/114397

**Authors:** Herty Liany, Jagath C. Rajapakse, R. Krishna Murthy Karuturi

**Author notes:** Email addresses: HL JCR RKMK.

## Abstract

**Background:** Differential co-expression signifies change in degree of co-expression of a set of genes among different biological conditions. It has been used to identify differential co-expression networks or interactomes. Many algorithms have been developed for single-factor differential co-expression analysis and applied in a variety of studies. However, in many studies, the samples are characterized by multiple factors such as genetic markers, clinical variables and treatments. No algorithm or methodology is available for multi-factor analysis of differential co-expression.

**Results:** We developed a novel formulation and a computationally efficient greedy search algorithm called MultiDCoX to perform multi-factor differential co-expression analysis of transcriptomic data. Simulated data analysis demonstrates that the algorithm can effectively elicit differentially co-expressed (DCX) gene sets and quantify the influence of each factor on co-expression. MultiDCoX analysis of a breast cancer dataset identified interesting biologically meaningful differentially coexpressed (DCX) gene sets along with genetic and clinical factors that influenced the respective differential co-expression.

**Conclusions:** MultiDCoX is a space and time efficient procedure to identify differentially co-expressed gene sets and successfully identify influence of individual factors on differential co-expression.

**Software:** R function will be available upon request.

## Background

Differential co-expression of a set of genes is the change in their degree of coexpression among two or more relevant biological conditions [12]. As illustrated in Figure 1, differentially co-expressed genes demonstrate a strong co-expression pattern among normal samples and no co-expression among disease samples. These genes may not be differentially expressed. Differential co-expression signifies loss of control of factor(s) over the respective downstream genes in a set of samples compared to the samples in which the gene set is co-expressed or variable influence of a factor in one set of samples over the other. This could also be due to a latent factor which had a significant influence on gene expression in a particular condition [44]. Since the proposal by Kostka & Spang [12], many algorithms have been proposed to identify *differentially co-expressed* (referred as DCX throughout the paper) gene sets and quantify differential co-expression. The algorithms can be classified based on two criteria: (1) method of identification of DCX gene sets (targeted, semi-targeted and untargeted); and (2) scoring method of differential co-expression (gene set scoring and gene-pair scoring).

**Figure 1.**
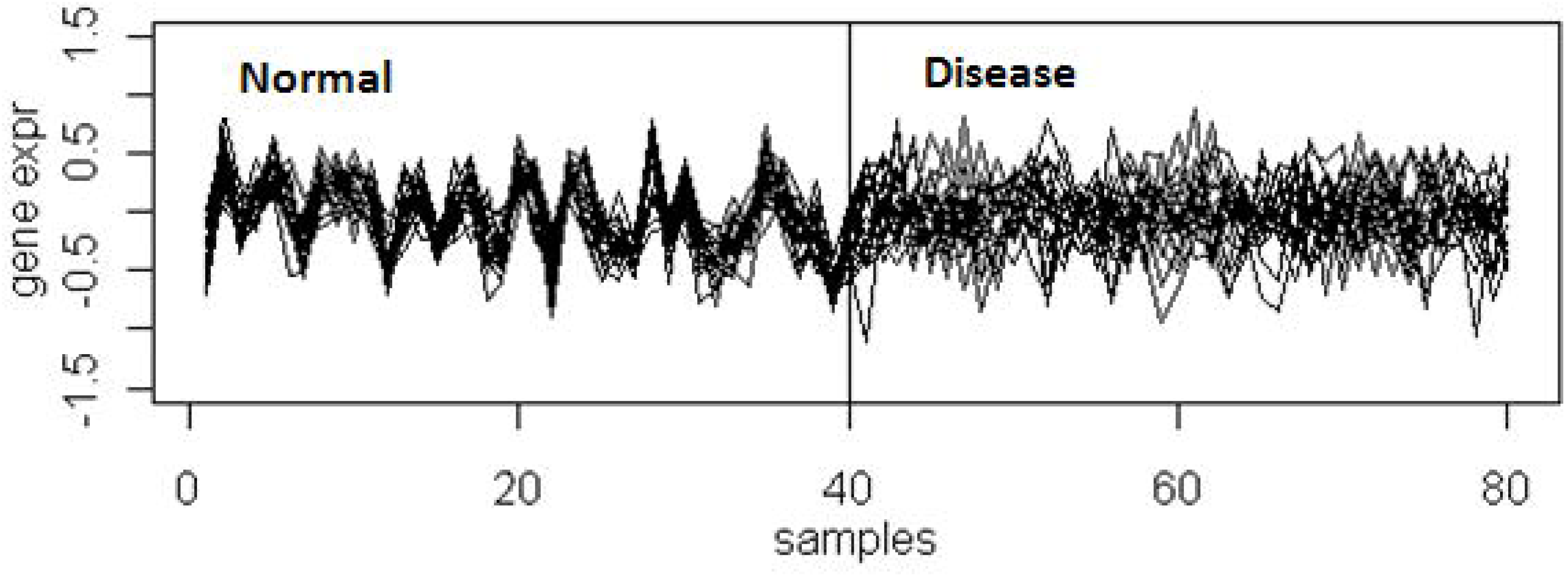
Differential Co-Expression. Geneset is co-expressed in normal samples but not in disease samples.

Based on the method of identification, similar to the one described by Tesson *et al.* [2], the algorithms can be classified into *targeted, semi-targeted* and *untargeted* algorithms. The *Targeted algorithms* [9] perform differential co-expression analysis on predefined sets of genes. The candidate gene sets may be obtained from public databases such as GO categories and KEGG pathways. They do not find novel sets of DCX gene sets. Another disadvantage of targeted methods is their reduced sensitivity if only a subset of the given gene set is differentially co-expressed as the DCX signal is diluted and the DCX geneset may not be identified. In addition, the DCX gene sets that are composed of genes of multiple biological processes or functions [44] may not be identified at all. The *semi-targeted algorithms* [2, 17, 24] work on the observation that the genes are co-expressed in one group of samples. Hence they perform clustering of genes in one set of samples, identify gene sets tightly co-expressed and test for their differential co-expression using the remaining group of samples. Although semi-targeted algorithms can identify novel gene sets, their applicability is limited to the co-expressed sets identified by the clustering algorithm. On the other hand, the *untargeted algorithms* [10, 12, 29] assume no prior candidate sets of genes and instead find the gene sets in an exploratory manner and therefore have a high potential to identify novel gene sets. The major drawback of untargeted approach is potentially high false discovery rate and large computational requirements.

The second aspect of DCX gene set identification algorithms is the methodology employed in scoring differential co-expression of a given gene set: (1) gene set scoring or set-wise method, and (2) gene pair scoring. In *gene set scoring,* all genes are considered in the scoring at once such as in the linear modelling used in Kostka&Spang [12] and Prieto *et al.* [10]. On the other hand, *gene-pair scoring,* as used in DiffFNs [29] and DCoX [2], computes differential correlation of each pair of genes in the gene set and summarizes them to obtain DCX score for the gene set. Gene pair scoring is intuitive and amenable to network like visualization and interpretation in single factor analysis settings. However, gene set scoring can be thought of as gene network of multiple cliques and cliques connected via common genes among all pairs of gene sets. The first few methods (e.g. Kostka&Spang [12] and Prieto *et al.* [10]) are untargeted set-wise methods, while DiffFNs [29] is an untargeted gene-pair scoring method. However, many later methods, including an early method (DCA [17]) are predominantly targeted or semi-targeted algorithms using gene pair scoring. Differential co-expression has been used in various disease studies and identified many interesting changed interactomes of genes among different disease conditions. DiffFNs [29], Differential co-expression [20], TSPG [39], and Topology-based cancer classification [22] were applied for the classification of tumor samples using interactome features identified using differential co-expression and shown good results over using individual gene features. The application of Ray and Zhang’s coexpression network using PCC and topological overlap on Alzheimer’s data helped identify gene sets whose co-expression changes in Alzheimer’s patients [27]. The multi-group time-course study on ageing [19] has identified gene sets whose coexpression is modulated by ageing. Application on data of *Shewanella oneidens* identified a network of transcriptional regulatory relationships between chemotaxis and electron transfer pathways [32]. Many other studies have also shown the significant utility of application of differential co-expression analysis [11, 14, 33, 36]. However, none of the existing algorithms allow direct multi-factor analysis of differential co-expression, i.e. deconvolving and quantifying the influence of different biological, environmental and clinical factors of relevance on the change in coexpression of gene sets. This is important as some phenotypes or biological outcomes are governed by multiple factors. In such a case, many single-factor differential coexpression analyses suffer from the same disadvantages of similar approach in differential expression analysis: it leads to multitude of tests, the interpretation of the identified gene sets may be cumbersome and misleading.

Multivariate differential co-expression analysis is important in many practical settings since each sample is characterized by many factors (a.k.a. co-factors) such as environmental variables, genetic markers, genotypes, phenotypes and treatments. For example, a lung cancer sample may be characterized by *EGFR* expression, smoking status of the patient, *KRAS* mutation and age [37]. Similarly, ageing of skin may depend on age, exposure to sun, race and sex [13]. Different environmental, genetic and clinical factors may modulate co-expression of a set of genes. Deconvolving and quantifying the effects of these factors on gene set’s co-expression and eliciting relevant regulatory pathways is an important task towards understanding the change in the cellular state and the underlying biology of interest.

Hence, we propose a very first methodology for such purpose called <u>Multi</u>-Factorial Analysis of <u>D</u>ifferential <u>Co</u>-e<u>X</u>pression or MultiDCoX, a gene set scoring based untargeted method. MultiDCoX performs greedy search for gene sets that maximize absolute coefficients of cofactors (as suggested in our earlier work [3]) while minimizing residuals for each geneset. Though the model used in MultiDCoX is restricted to ordinal and categorical valued factors, it is not a major limitation while dealing with real valued cofactors which can be discretized into reasonably small number of levels and be treated as ordinal variables. The analysis of several simulated datasets demonstrate that the algorithm can be used to reliably identify DCX gene sets and deconvolve and quantify the influence of multiple cofactors on the co-expression of a DCX geneset in the background of large set of non-DCX gene sets. The algorithm performs well even for genesets with weak signal-to-noise ratio. The analysis of a breast cancer gene expression dataset revealed interesting biologically meaningful DCX gene sets and their relationship with the relevant cofactors.

Furthermore, we have shown that the co-expression of CXCL13 is not only due to the Grade of the tumor as identified in [18], but also could be influenced by ER status. Similarly, MMP1 appears to play role in two different contexts defined by more than one co-factor. These together demonstrate the importance of multi-factor analysis.

## Methods

### MultiDCoX Formulation and Algorithm

MultiDCoX procedure consists of two major steps: (1) identifying DCX gene sets and obtaining respective DCX profiles; and (2) identifying covariates that influence differential co-expression of each DCX gene set. The formulation essential to carry out these two steps is as follows.

Let *E_im_* denote expression of gene *g_i_* in sample *S_m_.* The cofactor vector characterizing *S_m_* is denoted by *B_m_= (B_m1_, B_m2_, B_m3_, …, B_mk_)* where *B_mk_* is the value of *k*^th^ factor for *S_m_* which is either a binary or an ordinal variable. A categorical variable can be converted into as many binary variables as one less the number of categories of the factor.

We define a new variable *A_mn_(I)* to summarize co-expression of gene set *I* among sample pair *S_m_* and *S_n_* for which *B_m_* = *B_n_*, as

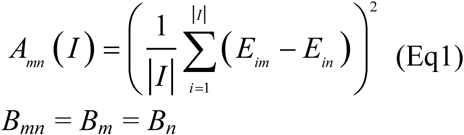

*A_mn_(I)* measures square of mean change of expression of all genes in *I* from *S_m_*to *S_n_.* Most of *A_mn_(I)’s* are expected to be non-zero among a group of samples in which I is co-expressed. On the other hand, if genes in *I* are not co-expressed in a group of samples then *A_mn_(I)’s* tend to be zero. This is illustrated in Figure 2.

**Figure 2.**
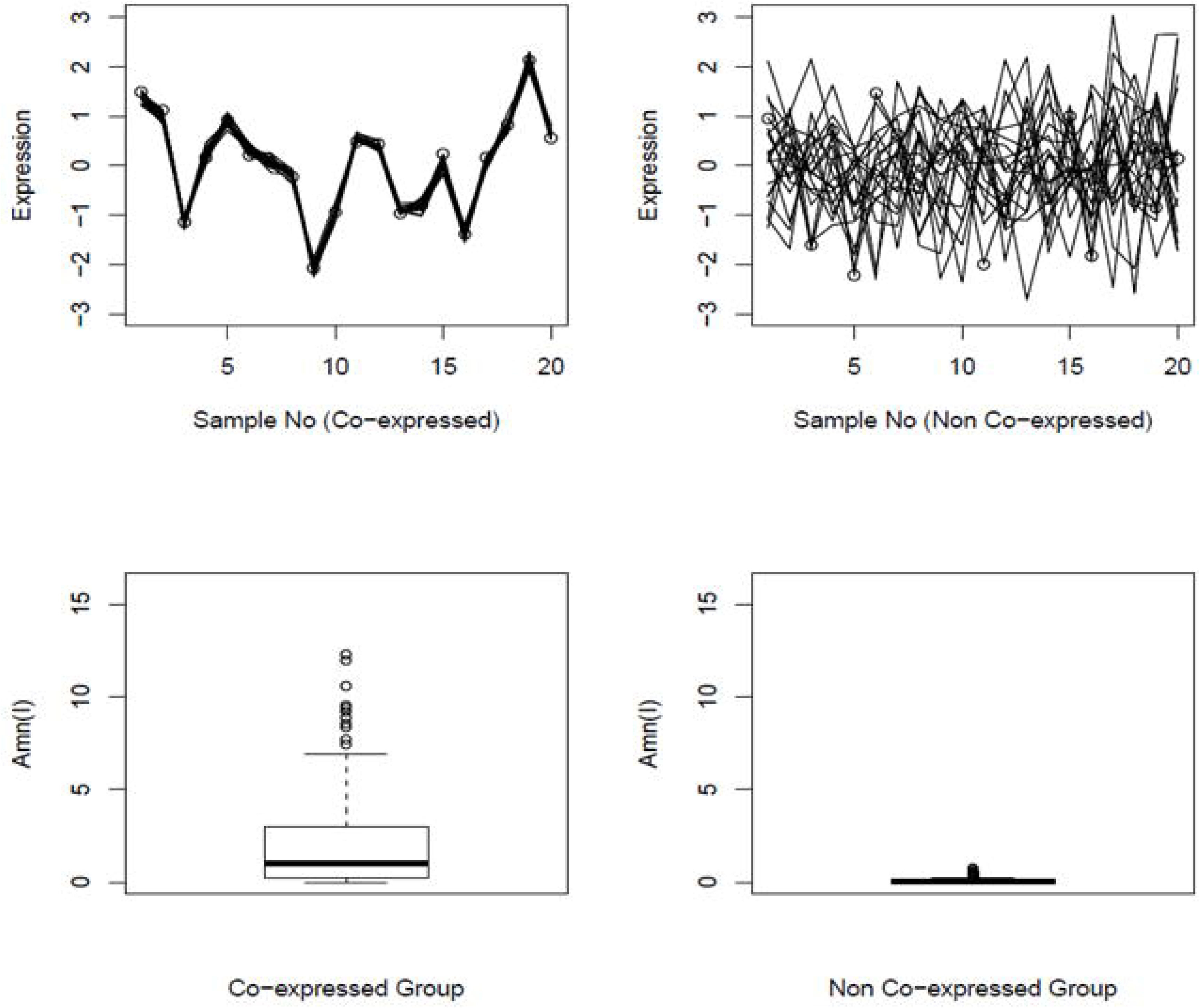
Illustration of A_mn_(I) for co-expression and non co-expression.

Once all *A_mn_(I)*s and *B_mn_*s are computed, we quantify the influence of the cofactors by fitting a linear model between *A_mn_(I)*s and *B_mn_s.* In other words, *A_mn_(I)s* are the instances of the response variable, *B_mn_*s form design matrix (B) and factors in the *B_mn_*s are explanatory variables or cofactors (F) i.e.

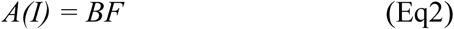

Where *A(I)* is the vector of *A_mn_(I)s, B* is matrix of *B_mn_s* and *F* is the vector of factors (attributes or covariates or cofactors) represented in *B_mn_s.* The coefficient vector obtained from the above modelling is called differential co-expression profile of the gene set *I*, denoted by *F(I).*

The linear modelling problem can be solved by standard functions (such as *lm())* available in R-package. But, identifying DCX gene sets is major computational task. The MultiDCoX algorithm identifies DCX gene sets by iteratively optimizing coefficient of a cofactor using the following procedure, see the flowchart in Figure 3 for the algorithm: (1) identifying significance threshold for cofactor coefficients; (2) choosing seed pairs of genes that demonstrate significant coefficient for the cofactor under consideration, i.e. the gene pairs may be differentially co-expressed for the cofactor; (3) expanding each chosen seed gene pair into a conservative multi-gene set by optimizing the respective coefficient; (4) augmenting the gene set to increase sensitivity or reduce false negatives while keeping the respective factor coefficient significant; and, (5) filtering out weak contributing genes from each gene set to increase specificity or reduce false positives. Each of these steps is explained in detail below.

**Figure 3.**
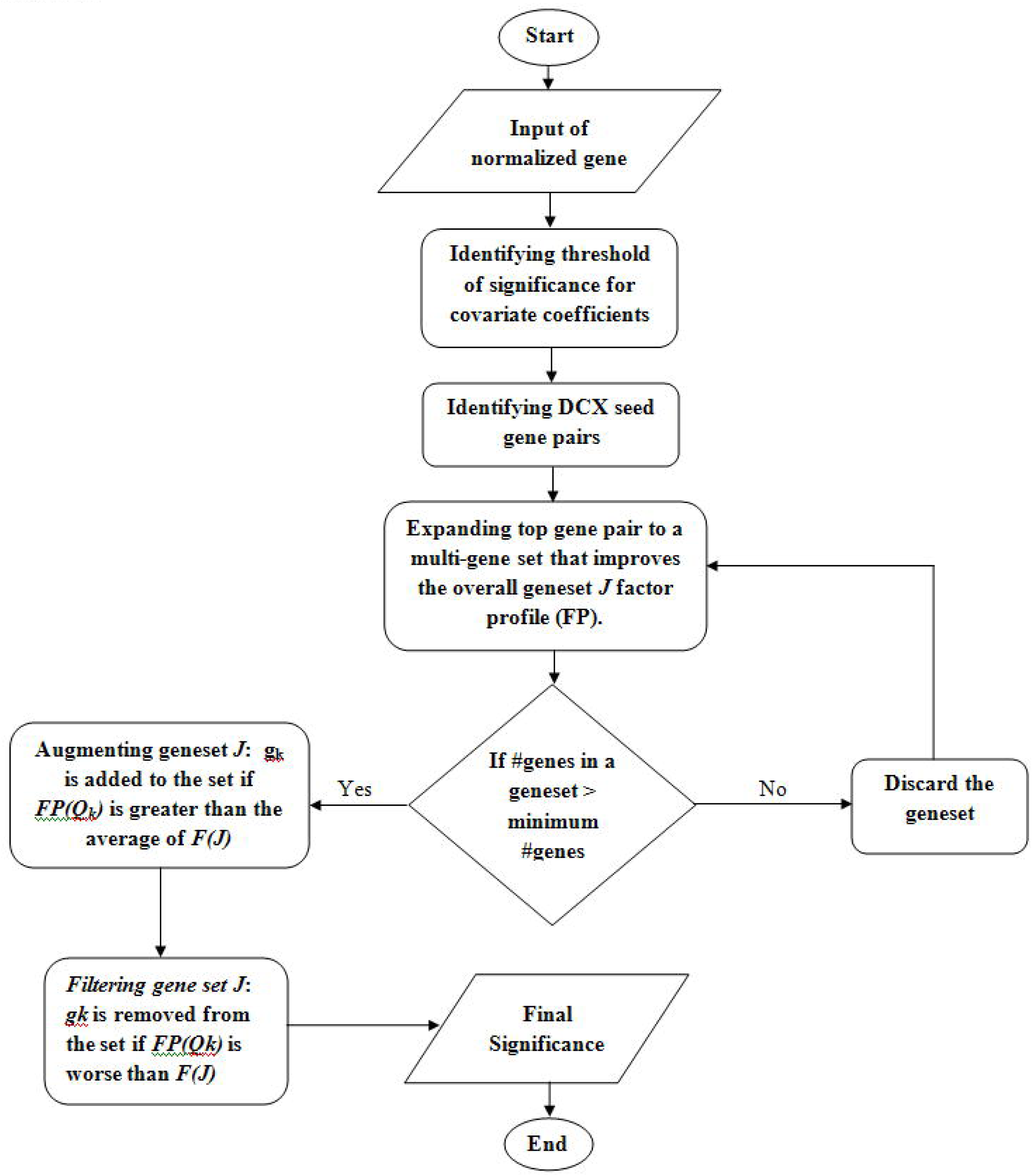
Flowchart of MultiDCoX algorithm.

#### 1. Identifying threshold of significance for cofactor coefficients

We generate the distribution of coefficients of the cofactors in *F* by random sampling of gene pairs: randomly sample large number of gene pairs, fit the linear model in Eq2 for each pair and obtain the coefficients in the linear models. Pool absolute values of coefficients of all factors of all gene pairs, and set half of the *m^t^*^h^ (*m*=10 in our experiments) highest value as absolute threshold for all cofactors. In other words,

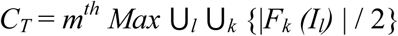

where*F_k_(I_Ɩ_)* is coefficient on gene set (a pair of genes in this case) *I_l_* for *k^th^* factor.

*T_oi_* is the threshold for cofactor ‘*i*’, derived from *C_T_* as follows

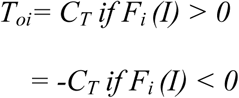

The division by 2 is necessary to avoid damagingly strict threshold and lay wider net at the beginning of the algorithm. *m*> 1 is required as some of the sampled gene pairs could belong to DCX gene sets which may overestimate the threshold and reduce sensitivity of the algorithm.

#### 2. Identifying DCX seed gene pairs

For each gene, search is performed throughout the dataset to find its partner gene whose pair can result in a linear model (Eq2) with at least one significant cofactor. A cofactor is considered to be significant if its linear model F-test p-value is < 0.01 and its coefficient is outside the range [*-C_T_, C_T_*]. If no partner gene could be found, then the gene will be filtered out from the dataset to improve the computational speed at later stages of the algorithm. We have implemented this step using the procedure: (a) batch application of *qr.coef()* in *R-package* which computes only linear model coefficients using one QR decomposition, (b) filter out gene pairs whose linear model coefficients are in the range [*-C_T_, C_T_*], (c) apply *lm()* on the gene pairs remaining after step ‘b’ to compute F-test p-values, and (d) further filter out gene pairs which do not meet requirements for the coefficient p-value. The batch application of *qr.coef()* is many folds faster than *lm().* We use similar strategy in the steps 3.A-3.C below to reduce computational requirements compared to the direct application of *lm().*

#### 3. Identifying DCXgene sets

We optimize coefficients of each significant cofactor for each gene pair in the direction, in positive or negative direction, depending on the sign of the coefficient i.e. if the coefficient is negative (positive) its minimized (maximized). To do so, for each factor, the steps 3.A-3.C are iterated until all seed pairs for which the factor is significant are exhausted from the seed pairs obtained in the step 2.

##### 3. A. Expanding top gene pair to a multi-gene set

We choose the gene pair whose constituent genes are not part of any of the multi-gene sets identified and whose linear model fit resulted in the highest coefficient for the factor of interest. It will be expanded to multi-gene set by adding genes that improve the coefficient of that factor in the direction of its coefficient for the gene pair. A sequential search is performed from first gene in the data to the last gene in the data (the order of the genes will be randomized before this sequential search). A gene is added to the set if it improved the coefficient of the factor under consideration i.e. the threshold to add a gene thereby the stringency increases as the search proceeds. The final set obtained at the end of this step is denoted by *J*. This step results in a most conservative DCX gene set. Factor profile *FP(J)* of *J* is defined as set of *(f_i_,h_i_)* pairs as follows:

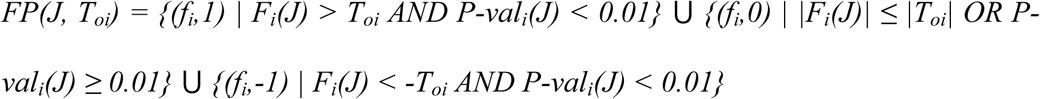

Where *f_i_* is factor *‘i’* and *h_i_* denotes whether it is positively *(h_i_=1)* or negatively *(h_i_=-* 1) significant or insignificant *(h_i_=0): F_i_(J)* is coefficient of factor *f_i_* for gene set *J P-val_i_(J)* is p-value of *F_i_(J)*

##### 3. B. Augmenting gene set J

As we tried to improve the coefficient of the factor for each addition of a gene in the expansion step (3.A), we may have missed many true positives which are not as strong constituents of *J*, but could be significant contributors. Therefore, we perform augmentation step to elicit some of the potential not-so strong constituents of *J* while preserving the factor profile of J. As the gene set identified in step (3.A) is most conservative, we set a new threshold *T_ni_(J)* or simply *T_ni_* for the coefficient *F_i_(J)* of each *f_i_* as

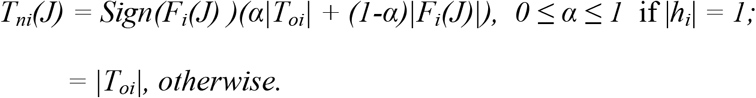

*T_ni_(J)* will be as stringent as *T_oi_* and at most equal to *F_i_(J)* which is the coefficient obtained at the end of step (3.A). Moreover, we define centroid *E_C_ (J) ={E_cm_(J)}* of *J* as

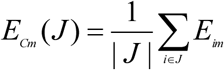

*E_C_(J)* is treated as a representative gene expression profile of *J* and find a gene sub set *K* such that each gene in *K, g_k_,* the pair *K_k_= (g_k_, E_C_(J))* satisfies the condition 

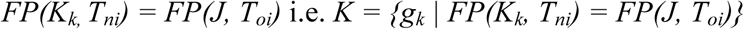

Then the augmented set *L = J&∪K* as new DCX gene set

##### 3. C. Filtering gene set L

The set *L* obtained after the step (3.B) may contain false positives which can be filtered out as follows: As in the augmentation step, we compute *E_cm_(L)* and evaluate each gene pair Q_k_ ∈ {(g_k_, *E_Cmn_(L))* | *g_k_* ∈ *L*} for *F(Q_k_). g_k_* is removed from the set if |*F_i_(Q_k_)|<F_i_(L)* for all *|h_i_|=1.* Then the final gene set *R = {g_k_ |g_k_*∈*L and F(Q_k_)* is better than *F(L)}. R* is the final set output for the run.

#### 4. Identifying cofactors significantly influencing DCX of each gene set

It is important to identify the factors influencing the DCX of a gene set to interpret the geneset and elicit underlying biology, i.e. *FP(R).* The F-test p-value obtained for each cofactor by the linear model fit (in Eq2) in the above procedure need to be further examined owing to the dependencies among the gene sets explored. Therefore, we mark a cofactor to be influential (|hi| =1) on co-expression of *R* if it satisfies the following two criteria:

a. *Effect size criterion*: We pool coefficients of all factors on all gene sets identified (denoted as *C_R_*) and examine their distribution. The valleys close to zero on either side of the central peak are chosen as the significance threshold *T_f+_ and T_f−_.,* see Figure 4. *F_i_(R)* is considered to be significant if it is >*T_f+_* or < *T_f−._* The underlying assumption is that not all factors influence all gene sets and the coefficients of the factors with no or little influence on certain gene sets will be suggestive of the distribution of the coefficients under null hypothesis.
b. *Permutation p-value criterion:* We permute the factor values of a DCX gene set, i.e. permute columns of B_mk_ matrix and fit the linear model in Eq2 for each gene set *R*. We repeat this procedure for a predefined number of iterations. A factor is said to be non-influential on the co-expression of the gene set under consideration if a minimum predefined fraction of permutations (0.01 in this paper) resulted in a fit in which the coefficient is better than *F_i_(R)* and its F-test p-value is better than the F-test p-value of the coefficient without permutation or 0.01 whichever is lower.

**Figure 4.**
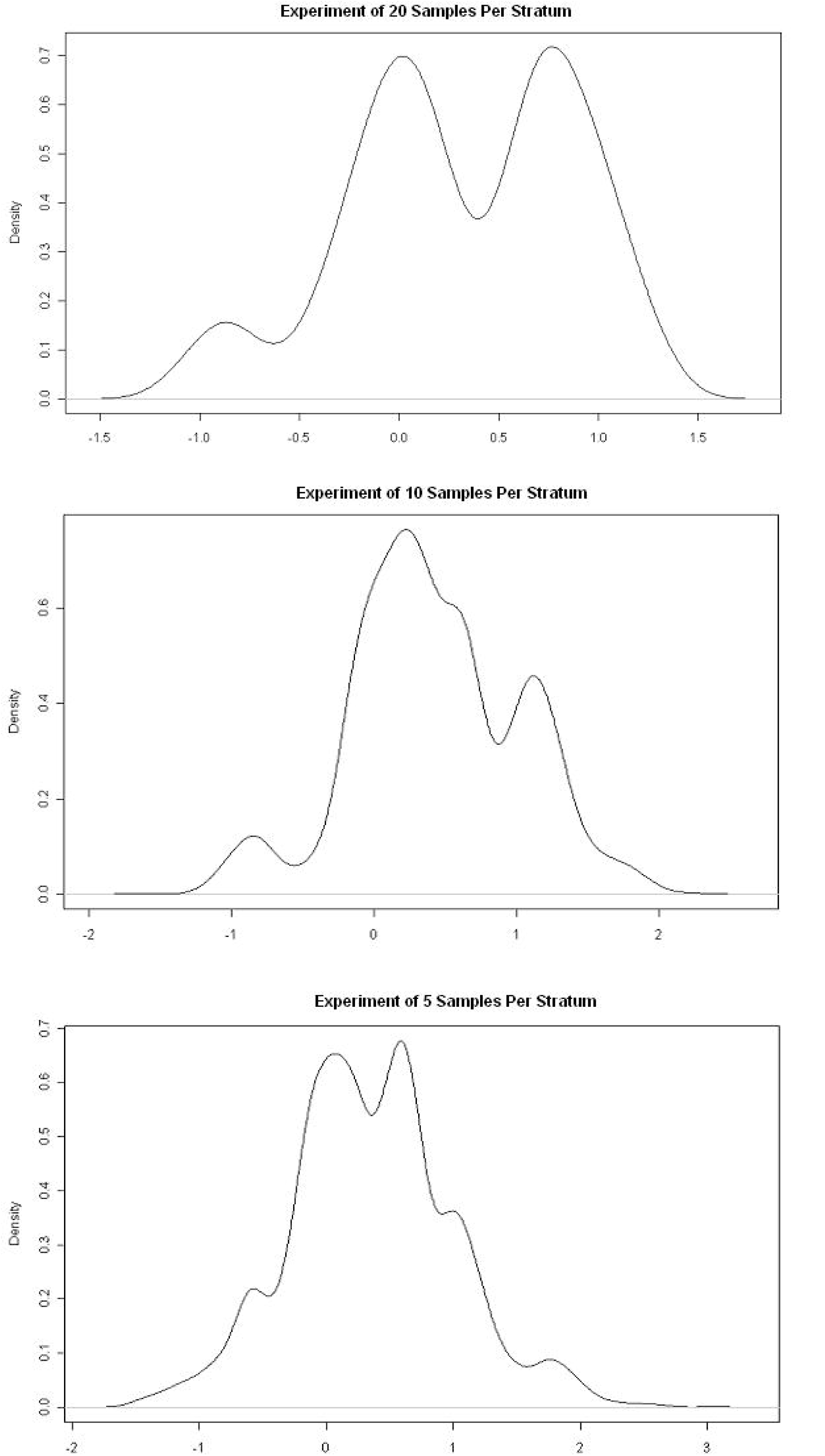
Density plots of all coefficients of the simulation data analysis by MultiDCoX for varying number of sample/stratum.

Finally, the gene sets with at least one significant cofactor and of predefined size (i.e. at least 6 genes in the set) will be output as DCX gene sets along with their factor profiles.

*Reducing computational and space requirements:* Computational and space requirements can be further reduced using the following strategies: (1) Filter out genes with no detectable signals among almost all samples and genes that demonstrate very little variance across the samples. This can result in modest reduction in space requirement and substantial reductions in computational requirement as the search procedure is at least of quadratic complexity if 50% of the genes are filtered out from the analysis; (2) Further reduction in computational time can be achieved in the step to identify seed gene pairs. Randomly split the genes into two halves and search for possible pairs where one belongs to one half and the other belongs to the other half, instead of all possible gene pairs. As many DCX gene sets are expected to be sufficiently large, >10 genes, the sampled set is expected to contain >2 genes from that set. This reduces computational time to find seed gene pairs by 2 fold. (3) Another possibility is to consider only a subset of sample pairs by randomly sampling a small fraction of (*m,n*)s for the linear model, it could be as small as 10% of all (*m,n*)s. We demonstrate the performance of the MultiDCoX even with such a minimal sampling. These three strategies put together with the optimization described in the step 2 of MultiDCoX can massively reduce the space and computational requirement by several folds and make the algorithm practically feasible.

## Results

### Simulation Results

To evaluate efficacy of MultiDCoX, we analysed simulated datasets of varying degrees of signal-to-noise ratio and sample size. Each simulated dataset consists of 50,000 probes as in a typical microarray and three factors of 12 stratums, sample sizes were chosen to be either 60 or 120 or 240 i.e. 5, 10 and 20 samples per stratum respectively. Two factors *B1* and *B2* were binary valued taking values from {-1, 1} and the other (B3) is an ordinal variable taking values from {-1, 0, 1}. Sample labels were randomly chosen for each factor and gene expression *(E_im_)* was simulated as described below:

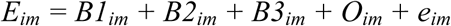

*B1_im_* = *B1_m_*~ N(0,1) if *S_m_* is in co-expressed group of *B1* and *g_i_* is in DCX gene set for the factor *B1,* 0 otherwise. Similar interpretation holds for the remaining factors, B2 and B3, too. *O_im_= O_m_*~ N(0,1) indicates co-expression over all samples if *g_i_* belongs to set of genes co-expressed across all samples irrespective of the factor values. *e_im_~ Ν(0,σ*^2^) is noise term and *σ*^2^ is the extant of noise in the data.

We simulated 20 genes which show co-expression for *B1_m_=1* and *B2_m_=1,* 20 genes co-expressed for *B1_m_= −1* only, and another 20 genes with *O_i_= 1* only. With this we have two sets of negative controls: large number of genes with no co-expression and a set of genes co-expressed across all samples. Ideally, a DCX gene set identification algorithm should be able to discriminate the first two sets of genes from the two control (negative) sets. Furthermore, we have tested our MultiDCoX for three different values of *σ ∈ {0.2, 0.5, 0.8}* i.e. from low noise to the noise comparable to the signal. We carried out 10 simulations for each choice of σ.

The simulation results are summarized in the panel of plots in Figure 5: plots of average numbers of false positives (FPs) and false negatives (FNs) over 10 independent runs for each choice of *σ* and sample size for both DCX gene sets along with the globally co-expressed gene set. MultiDCoX performed well in terms of both false positives and false negatives for low to medium values of *σ*. Moreover, the algorithm exhibited reasonable performance even at the noise (*σ*) comparable to the signal (i.e. *σ* = 0.8). The simulation results also demonstrate that MultiDCoX is sensitive even at small sample size for low to medium noise level. The failure rate of identifying gene sets and their profiles are dependent not only on the sample size and noise level, but also on the type of set identified. A single factor influenced set has better chance of identifying the right profile but poor chance of being identified at low sample size and higher noise level. On the other hand, the set influenced by 2 factors has higher chance of being identified, but poorer chance of being identified with correct profile at low sample size and higher noise level. The effect of noise on FNR also depended on the number of factors influencing the DCX gene set. However, FDR is less dependent on both noise level and the number of factors influencing coexpression. The number of simulations that identified false gene sets increased with increased noise and reduced sample size. It is the lowest for 5 samples/stratum and high noise (σ = 0.8). The computational time for MultiDCoX analysis, to optimize each cofactor in both directions (maximization and minimization), was ~12-15 hours for one simulated data of 240 samples using 1 node of a typical HPC cluster.

**Figure 5.**
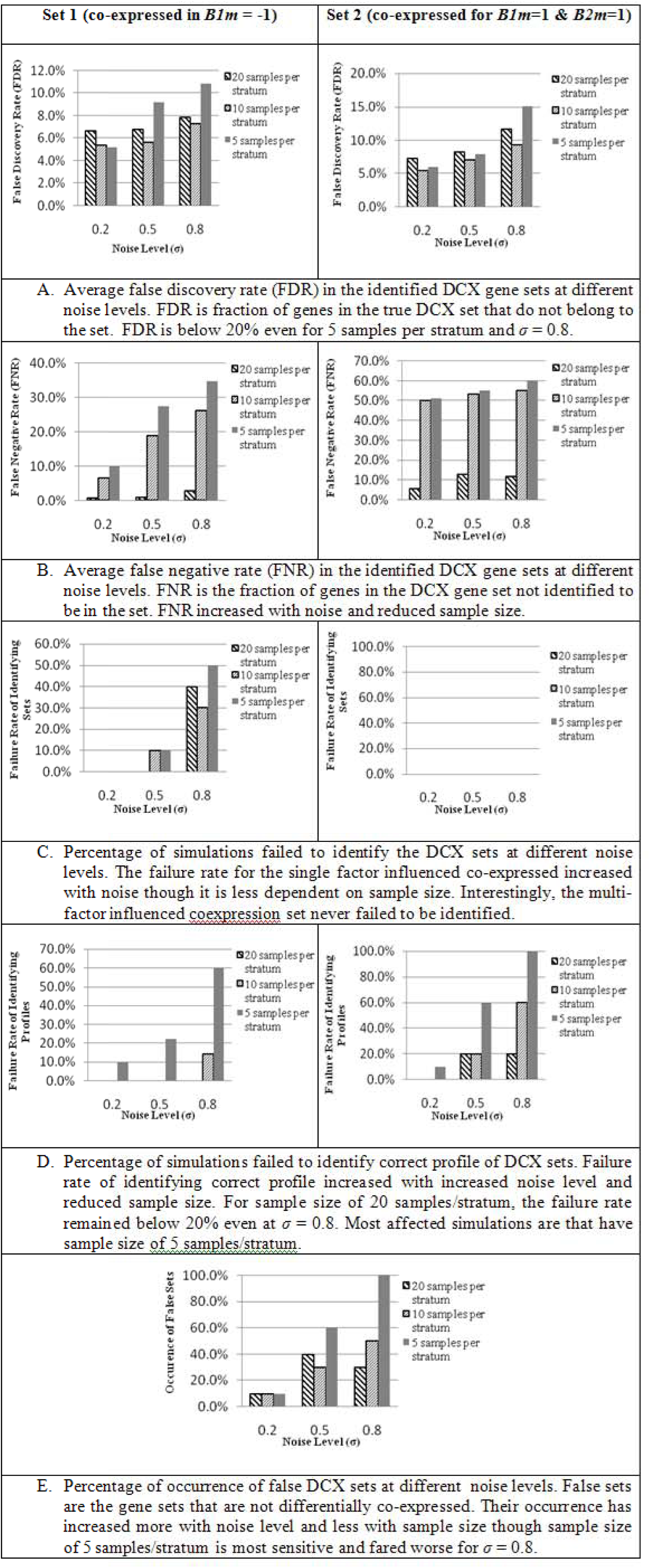
Simulation results. The simulations were carried out for 5 samples/stratum, 10 samples/stratum and 20 samples/stratum. Set 1 represents gene set simulated to be co-expressed only in samples *B1_m_= −1,* while Set 2 represents gene set simulated to be co-expressed for *B1_m_=1* and *B2_m_=1.*

### MultiDCoX Analysis of Breast Cancer Data

We analyzed a breast cancer gene expression data published by Miller *et al.* [25]. It contains expression profiles of 258 breast cancer patients on U133A and U133B Affymetrix arrays i.e. ~44,000 probes. The tumors were annotated for their oestrogen receptor (ER) status (1 for recognizable level of ER or *ER+,* −1 otherwise or *ER-),* p53 mutational status (1 for mutation or *p53+,* and −1 for wild type or *p53-)* and grade of tumor (-1 for grade 1, 0 for grade 2 and 1 for grade 3). ER and p53 status are the important markers used to guide treatment and prognosis of breast cancer patients. Hence it is important to study the gene sets regulated and thereby co-expressed by these factors while accounting for the effect of the tumor status as indicated by its grade and strong association between these three factors. For example, p53-mutant tumors are typically of higher grade (grades 2 or 3) tumors with correlation ~ 63% [40] and ER-positive tumors are typically low grade (grade 1) tumors [41]. In the presence of these correlations among the covariates, it is important to identify and quantify their effects on co-expression of gene sets. We have applied *MultiDCoX* on this dataset using ER status, p53 mutational status and tumor grade as cofactors. We discuss a few DCX sets here and the remaining DCX gene sets are given in the ***Additional File 1*** *(UppsalaBCResults.xlsx).*

#### Co-expression of ER pathway and the genes associated with relevant processes is modulated in p53 mutated tumors

A DCX gene set and the linear model fit is shown in Table 1A. The set is co-expressed only in p53 mutant tumors. The co-expression plot of p53 mutant tumors is shown in Figure 6.

**Figure 6.**
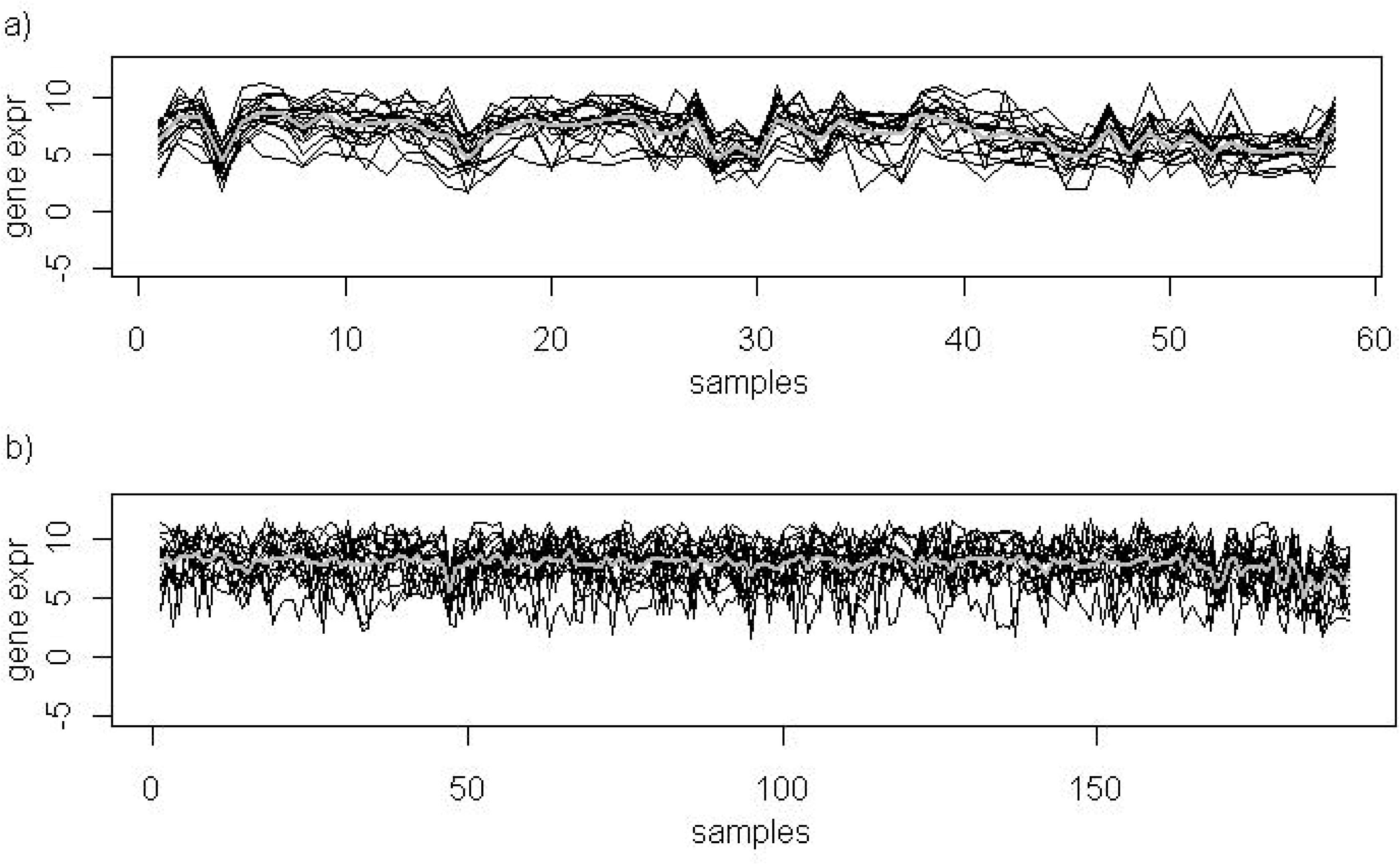
The co-expression plot of set 1 (Table 1A) in *p53+* tumors in breast cancer data. a. Co-expression of geneset 1 (18 genes) across p53 mutant tumor (p53+) samples; gray color line indicates mean expression value of geneset 1.
b. The geneset 1 showed no co-expression in p53 wild-type samples (p53-); gray color line indicates mean expression value of geneset 1.

**Table 1A.** The table shows the MultiDCoX model fit over three factors (ER, p53 and Grade). A gene set identified is differentially co-expressed in p53+ with F-test p-value = 2.75 x 10^-231^ and coefficients value = 1.137. Only p53 covariate is significant and its coefficient is positive which means the co-expression of the set occurs only in p53 mutant tumors only.

| Co-expression | Genes | Total_Genes | ER coefficient | ER pvalue | p53 coefficient | p53 pvalue | Grade coefficient | Grade pvalue |
| --- | --- | --- | --- | --- | --- | --- | --- | --- |
| p53+<br>(p53 mutant) | MKX, GFRA1, GATA3, SPDEF, GAMT, TOX3, FOXA1, AGR3, ESR1, SDR16C5, PIP, CYP2B7P1, SYTL5, REEP6, AGR2, ANKRD30A, CA12, SCGB2A1 | 18 | 0.087 | 0.114 | **1.137** | **2.75E-231** | -0.063 | 0.028 |

**Table 1B.** ER dependent differential expression, ER binding sites and p53 binding sites for the geneset in Table 1A.

| No. | Gene | ER (DE) | ER Binding Site | p53 Binding Site | Gene Description |
| --- | --- | --- | --- | --- | --- |
| 1. | GFRA1 | **Yes (up)** | **Yes(dist=58.5kb)** | No | TGF-beta related neurotrophic factor receptor |
| 2. | FOXA1 | No | **Yes(dist=4.798kb)** | No | Forkhead box protein A1 |
| 3. | GATA3 | No | **Yes(dist=30.338kb)** | **Yes** | GATA binding protein 3 |
| 4. | SPDEF | No | **Yes(dist=1.159kb)** | No | SAM pointed domain containing ets transcription factor |
| 5. | ESR1 | **Yes (up)** | **Yes(dist=32.241kb)** | **Yes** | Estrogen receptor 1 |
| 6. | GAMT | No | dist > 100kb | **Yes** | guanidinoacetate N-methyltransferase |
| 7. | TOX3 | No | dist > 100kb | No | TOX high mobility group box family member 3 |
| 8. | AGR3 | **Yes (up)** | **Yes(dist=54.069kb)** | No | anterior gradient 3 homolog (Xenopus laevis) |
| 9. | SDR16C5 | No | dist > 100kb | No | Short-chain dehydrogenase/reductase family 16C member 5 |
| 10. | PIP | No | dist > 100kb | No | prolactin-induced protein |
| 11. | CYP2B7P1 | No | dist > 100kb | No | cytochrome P450, family 2, subfamily B, polypeptide 7 pseudogene 1 |
| 12. | SYTL5 | **Yes (up)** | Yes(dist=94.215kb) | No | synaptotagmin-like protein 5 |
| 13. | MKX | No | **Yes(dist=35.211kb)** | No | mohawk homeobox |
| 14. | REEP6 | No | dist > 100kb | No | receptor accessory protein 6 |
| 15. | AGR2 | **Yes (up)** | **Yes(dist=2.154kb)** | No | anterior gradient 2 homolog (Xenopus laevis) |
| 16. | ANKRD30A | No | dist > 100kb | No | ankyrin repeat domain 30A |
| 17. | CA12 | **Yes (up)** | **Yes(dist=56.695kb)** | **Yes** | Carbonate dehydratase XII |
| 18. | SCGB2A1 | No | dist > 100kb | No | secretoglobin, family 2A, member 1 |

The set includes *ESR1* (which encodes *ERα)* and its co-factor *GATA3* and pioneering factor *FOXA1* [23] along with ER downstream targets *CA12, SPDEF* and *AGR2.* We retrieved a total of 1349 p53 binding sites’ associated genes data from Botcheva K *et al.* [1], and Wei CL *et al.* [5]. p53 binding sites are reported to be close to the promoters of *ESR1* [30] as well as *GATA3.* Furthermore, *GATA3* appears to bind to *FOXA1* which is a pioneering factor of ER [35]. Our finding reinforces the observations made by Rasti*et al.* [30] that different p53 mutations may have varying effect on the expression of *ESR1* gene, it’s co-factor *GATA3,* pioneering factor *FOXA1* and SAM-dependent Mythyltransferase & *p53* interacting *GAMT* which could have resulted in the differential co-expression of the ER pathway. In addition, comodulation of chromatin structure alternating &*ER* promoter stimulating *TOX3* and Protein transfer associated REEP6 appears to be required to modulate ER pathway by p53.

#### Genes co-expressed with BRCA2 in ER-negative tumors are associated with Her2-neu status

Another gene set of interest is co-expressed in ER-negative tumors only and its details are given in Tables 2A and 2B. The co-expression plot of the gene set in ER-negative tumors is shown in Figure 7. The gene set includes tumor suppressor gene *BRCA2.* We have investigated ER binding sites published by Carroll *et al.* [4] and Lin *et al.* [5] for ER binding sites close (within ±35Kb from TSS) to these genes. The ~4800 binding sites mapped to ~1500 genes. Significantly, 10 of the 21 genes in this DCX gene set have ER binding sites mapped to them which is statistically significant at F-test p-value < 0.01. Interestingly, most of these genes have not been identified to be ER regulated in the earlier studies using differential expression methodologies, possibly owing to the complexity of regulatory mechanisms. However, many of these genes are down regulated in ER-negative tumors. Testing for association of expression of this set with Her2-neu status revealed that higher expression in ER-negative tumors is associated with Her2-neu positivity which must have led to co-expression in ER-tumors. Odds ratio of such an association is 18 which is much higher than that of ER positive tumors (OR = 4).

**Figure 7.**
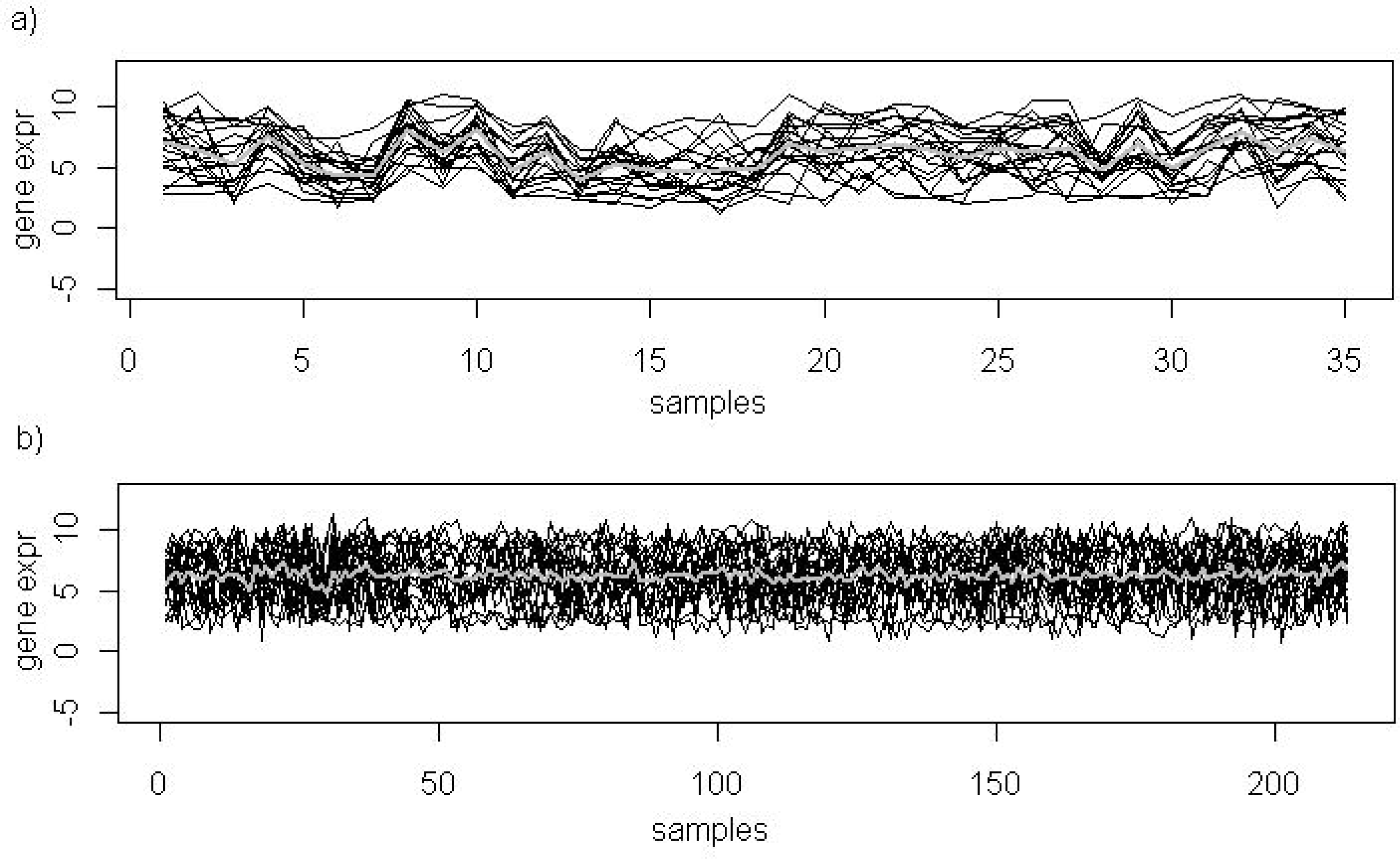
The co-expression plot of set 2 (Table 2A) tumors in breast cancer data. a. Co-expression of geneset 2 (21 genes) in ER-negative tumor samples; gray color line indicates sample-wise mean expression value of it.
b. The geneset 2 showed no co-expression in ER-positive tumor samples; gray line indicates mean expression value.

**Table 2A.**
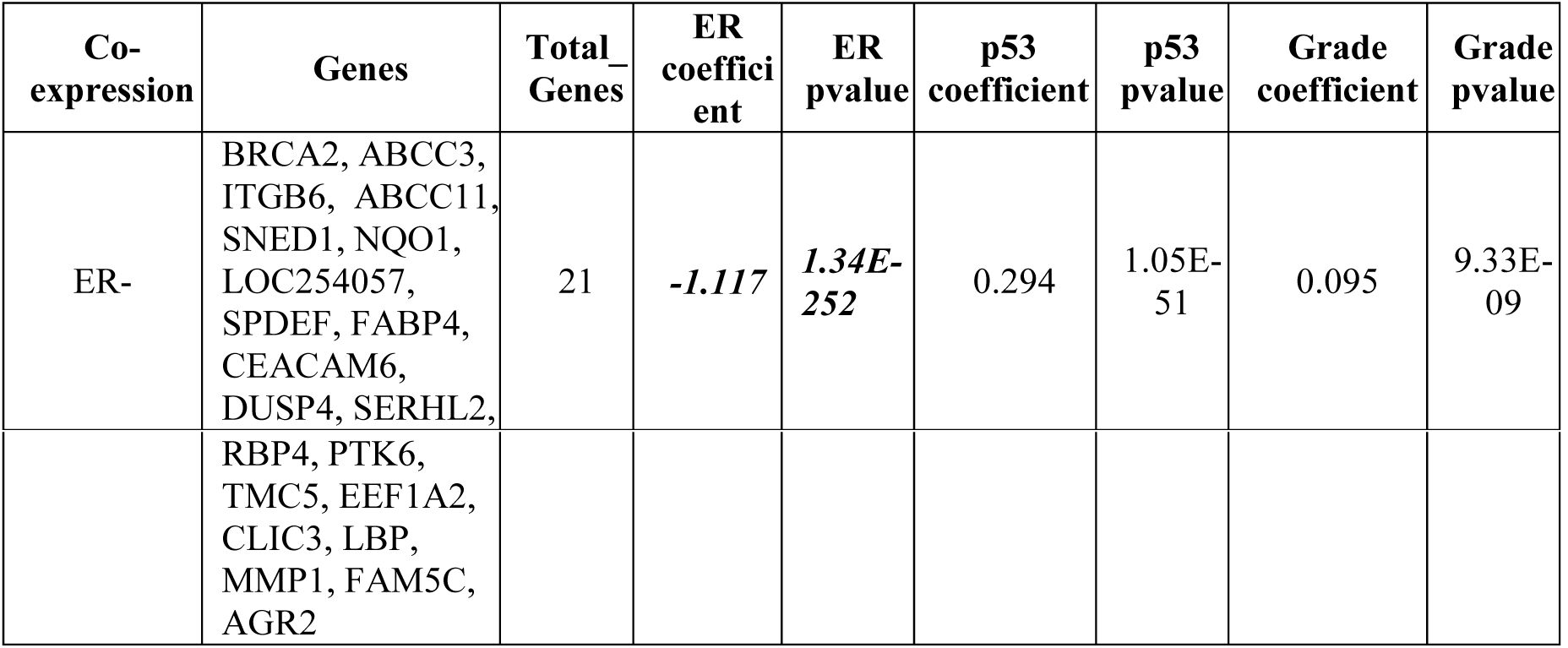
The table shows the MultiDCoX model fit over three factors (ER, p53 and Grade). A gene set identified is differentially co-expressed in ER-negative covariate with F-test p-value = 1.34 x 10^-252^ and coefficients value = −1.117. Only ER covariate is significant and its coefficient is negative which means the co-expression of the set occurs in ER-negative tumors only.

| Co-expression | Genes | Total Genes | ER coefficient | ER pvalue | p53 coefficient | p53 pvalue | Grade coefficient | Grade pvalue |
| --- | --- | --- | --- | --- | --- | --- | --- | --- |
| ER- | BRCA2, ABCC3, ITGB6, ABCC11, SNED1, NQO1, LOC254057, SPDEF, FABP4, CEACAM6, DUSP4, SERHL2, | 21 | **-1.117** | **1.34E-252** | 0.294 | 1.05E-51 | 0.095 | 9.33E-09 |
|  | RBP4, PTK6,<br>TMC5, EEF1A2,<br>CLIC3, LBP,<br>MMP1, FAM5C,<br>AGR2 |  |  |  |  |  |  |  |

**Table 2B.** ER dependent differential expression, ER binding sites and p53 binding sites for the gene set in Table 2A.

| No. | Gene | ER (DE) | ER Binding Site | Gene Description |
| --- | --- | --- | --- | --- |
| 1. | BRCA2 | **Yes (up)** | dist > 100kb | breast cancer 2, early onset |
| 2. | ABCC3 | *Yes (down)* | **Yes(dist=20.96kb)** | ATP-binding cassette, sub-family C (CFTR/MRP), member 3 |
| 3. | ITGB6 | *Yes (down)* | dist > 100kb | integrin, beta 6 |
| 4. | ABCC11 | No | **Yes(dist=68.96kb)** | ATP-binding cassette, sub-family C (CFTR/MRP), member 11 |
| 5. | SNED1 | No | **Yes(dist=94.62kb)** | Insulin-responsive sequence DNA-binding protein 1 |
| 6. | NQO1 | *Yes (down)* | **Yes(dist=32.63kb)** | NAD(P)H dehydrogenase, quinone 1 |
| 7. | LOC254057 | No | NA | uncharacterized LOC254057 |
| 8. | SPDEF | No | **Yes(dist=1.159kb)** | SAM pointed domain containing ets transcription factor |
| 9. | FABP4 | No | **Yes(dist=1.159kb)** | fatty acid binding protein 4, adipocyte |
| 10. | CEACAM6 | *Yes (down)* | **Yes(dist=19.05kb)** | carcinoembryonic antigen-related cell adhesion molecule 6 |
| 11. | DUSP4 | No | **Yes(dist=19.138kb)** | dual specificity phosphatase 4 |
| 12. | SERHL2 | No | **Yes(dist=32.63kb)** | serine hydrolase-like 2 |
| 13. | RBP4 | No | **Yes(dist=20.489kb)** | retinol binding protein 4, plasma |
| 14. | PTK6 | *Yes (down)* | dist > 100kb | PTK6 protein tyrosine kinase 6 |
| 15. | TMC5 | No | dist > 100kb | transmembrane channel-like 5 |
| 16. | EEF1A2 | No | dist > 100kb | eukaryotic translation elongation factor 1 alpha 2 |
| 17. | CLIC3 | *Yes (down)* | **Yes(dist=0.317kb)** | chloride intracellular channel 3 |
| 18. | LBP | No | dist > 100kb | lipopolysaccharide binding protein |
| 19. | MMP1 | No | dist > 100kb | matrix metalloproteinase 1 (interstitial collagenase) |
| 20. | FAM5C | No | dist > 100kb | family with sequence similarity 5, member C |
| 21. | AGR2 | **Yes (up)** | **Yes(dist=2.154kb)** | anterior gradient 2 homolog (Xenopus laevis) |

### DCX of CXCL13 is modulated by Grade and ER status too

Analysis of Grade1 and Grade3 tumors using GGMs [18] helped identify *CXCL13* in breast cancer as hub gene. It emerged as one of the hub genes in our analysis too, contributing to multiple DCX gene sets (see **Additional File 1**, sheet*: maxGrade*). Although they are significant for Grade, they are significant for ER status too. It shows that CXCL13’s differential co-expression appears to be influenced by ER status, in addition to Grade. This couldn’t be identified in the previous study as it was restricted to univariate (Grade) analysis.

### DCX of MMP1 is modulated by factor subspace associated with poor survival

MMP1 is another gene we have examined whose family of genes are associated with poor survival [43]. MMP1 is co-expressed among tumors which are P53+ (mutant) and ER-negative or hi-grade tumors which are ER-postive (see ***Additional File 1***, sheets:*maxP53, maxGrade and minER).* Both these categories are known to be associated with poor survival of patients. This couldn’t have been revealed in a single factor analyses.

### Functional analysis of DCX profiles

To elucidate the biological function of different DCX profiles (ER+, ER-& p53+, etc.), we pooled all genes from gene sets of same DCX profile and used DAVID functional annotation tool [16] to identify GO terms, protein domains, tissues of expression and pathways enriched, the results are tabulated in Table 3. It shows a clear distinction of GO functional categories, pathways enriched and tissues of expression between different DCX profiles. For example there is distinctive protein domains between ER-positive and ER-negative DCX profiles, whereby ER-positive’s protein domain involves more in Immunoglobulin/major histocompatibility complex while ER-negative involves in epidermal growth factor (EGF) extracellular domain of membrane-bound proteins. Also, both ER-negative and ER-positive covariates are associated with different pathway categories: ER-positive’s pathway involves more in synaptic transmission, neuroactive ligand-receptor interaction while ER-negative is associated with hormone (steroid, androgen and estrogen) metabolism, drug, starch and sucrose metabolism. The same phenomena can be observed for p53-mutant versus p53-negative associated genes.

**Table 3.** Functional analysis of co-expression in different covariates. GO (Gene ontology) summary shows gene sets, GO terms enriched and the influence of different cofactors. For each covariate, the sets with positive coefficient are pooled and analyzed for GO and pathway enrichment, it is repeated for the sets with negative coefficient as well.

| Significant Covariates | Biological Process | Protein Domain | Tissue Expression | Pathways (KEGG/BIOCARTA/PANTHER) |
| --- | --- | --- | --- | --- |
| ER+ | cell adhesion, biological adhesion, cell-cell adhesion (Enrichment Score: 4.15) | Immunoglobulin C1-set, IGc1, Immunoglobulin/major histocompatibility complex, conserved site, Immunoglobulin-like, Immunoglobulin V-set, subgroup, IGv, Immunoglobulin-like fold, Immunoglobulin V-set, immunoglobulin V region (Enrichment Score: 3.27) | Smooth Muscle_3rd, WHOLE BLOOD_3rd, TONGUE_3rd, BM-CD105+Endothelial_3rd (Enrichment Score: 4.22) | Synaptic Transmission, Ionotropic glutamate receptor pathway, Metabotropic glutamate receptor group III pathway, Neuroactive ligand-receptor interaction, Amyotrophic lateral sclerosis (ALS) (Enrichment Score: 1.89) |
| ER- | extracellular region, extracellular region part, extracellular space (Enrichment Score: 15.33) | EGF-like region, conserved site, EGF-extracellular, EGF-like, EGF (Enrichment Score: 1.98) | Uterus_3rd, Cingulate Cortex_3rd, WHOLE BLOOD_3rd, BM-CD105+Endothelial_3rd, bone marrow_3rd, Smooth Muscle_3rd, TONGUE_3rd (Enrichment Score: 22.75) | Drug metabolism, Androgen and estrogen metabolism, Metabolism of xenobiotics by cytochrome P450, Steroid hormone biosynthesis, Retinol metabolism, Starch and sucrose metabolism, Ascorbate and aldarate metabolism, Pentose and glucuronate interconversions (Enrichment Score: 2.47) |
| P53+ | extracellular region, extracellular region part, extracellular space (Enrichment Score: 6.32) | Immunoglobulin C1-set, IGc1, Immunoglobulin/major histocompatibility complex, conserved site, Immunoglobulin-like, Immunoglobulin-like fold, IGv, Immunoglobulin V-set (Enrichment Score: 2.51) | Smooth Muscle_3rd, Uterus_3rd, WHOLE BLOOD_3rd, TONGUE_3rd RT, BM-CD105+Endothelial_3rd (Enrichment: 8.23) | Drug metabolism RT, Metabolism of xenobiotics by cytochrome P450 RT, Retinol metabolism (Enrichment Score: 1.11) |
| P53- | membrane-bounded vesicle, vesicle, cytoplasmic membrane-bounded vesicle, cytoplasmic vesicle, secretory granule, synapse (Enrichment Score: 1.36) | Cytochrome P450, Cytochrome P450 E-class-group I, Cytochrome P450, conserved site (Enrichment Score: 1.79) | spinalcord_3rd, WHOLE BLOOD_3rd, Smooth Muscle_3rd, TONGUE_3rd, BM-CD105+Endothelial_3rd (Enrichment Score: 2.87) | None |
| Gr+ | extracellular region, extracellular region part, extracellular space (Enrichment Score: 13.29) | Small chemokine, interleukin-8-like, C-X-C, conserved site, CXC chemokine, C-X-C/Interleukin 8, small inducible chemokine (Enrichment Score: 4.07) | uncharacterized tissue_uncharacterized histology_3rd (Enrichment Score: 1.96) | Chemokine_families, Chemokine signaling pathway, Cytokine-cytokine receptor interaction (Enrichment Score: 3.17) |
| Gr- | extracellular region, extracellular region part, extracellular space (Enrichment Score: 1.85) | Protein-tyrosine phosphatase, Dual-specific/protein-tyrosine phosphatase, Protein-tyrosine phosphatase, active site (Enrichment Score: 1.01) | TemporalLobe_3rd, WHOLE BLOOD_3rd, TONGUE_3rd, Smooth Muscle_3rd, BM-CD105+Endothelial_3rd, bone marrow_3rd, Uteru s_3rd,Cingulate<br>Cortex_3rd<br>(Enrichment<br>Score: 3.5) | None |
| ER- &<br>P53+ | extracellular region,<br>extracellular region<br>part, extracellular<br>space (Enrichment<br>Score: 3.08) | cytoskeletal<br>keratin,Filament,Intermedi<br>ate filament protein-<br>conserved site,Keratin-<br>type I (Enrichment Score:<br>3.48) | Uterus_3rd,Cingul<br>ate<br>Cortex_3rd,bone<br>marrow_3rd<br>(Enrichment<br>Score: 2.08) | None |
| ER+ & Gr+ | cell-cell signaling,<br>transmission of nerve<br>impulse, synaptic<br>transmission<br>(Enrichment Score:<br>2.49) | Immunoglobulin C1-<br>set,IgC1,Immunoglobulin/<br>major histocompatibility<br>complex, conserved site<br>,Immunoglobulin-<br>like,Immunoglobulin V-<br>set,<br>subgroup,IGv,Immunoglo<br>bulin-like<br>fold,Immunoglobulin V-<br>set,immunoglobulin V<br>region (Enrichment Score:<br>5.38) | Trigeminal<br>Ganglion_3rd,skin<br>_3rd,Trachea_3rd<br>(Enrichment<br>Score: 7.93) | Intestinal immune<br>network for IgA<br>production,Type I<br>diabetes mellitus,Cell<br>adhesion molecules<br>(CAMs),Asthma,Allograft<br>rejection,Graft-versus-<br>host disease,Autoimmune<br>thyroid disease,Viral<br>myocarditis,Antigen<br>processing and<br>presentation,Systemic<br>lupus erythematosus<br>(Enrichment Score: 1.32) |

## Discussion

MultiDCoX is a space and time efficient algorithm which successfully elicits quantitative influence of cofactors on co-expression of gene sets. It required only 12 hours of computation on a typical HPC node to identify DCX gene sets for each factor for a dataset of 240 samples and ~44000 probes. The simulation results demonstrated that MultiDCoX has tolerable false discovery rates even at 5 samples/stratum and noise (σ) of 0.8. However, false negative rate (FNR) was affected by both sample size and noise level. As expected, FNR is very low for large sample size (20 samples per stratum) and low noise level (σ = 0.2). Interestingly, both FDR and FNR did not greatly depended on the type of the gene set to be discovered, or whether it is influenced by single factor or multi-factors. The discovery of a gene set whose DCX is driven by two cofactors is less affected by noise and sample size than the gene sets influenced by a single cofactor. On the other hand, the set influenced by 2-cofactors has higher likelihood of arriving at the wrong profile compared to that of a 1-cofactor driven DCX. Occurrence of false DCX sets increased with increasing noise level, it is pronounced more for small sample size case. This is a major issue to be addressed in the future improvements over the current version of MultiDCoX. Moreover, the performance of the algorithm needs to be studied for varying parameters’ settings and further reductions in computational time. It is possible to reduce the computational time by 2 fold by filtering out 50% of probes of low variance in expression. Though we have not used this strategy as we needed to study its impact on the discovery and profiling of DCX gene sets, the current implementation could complete the analysis within half a day of computing for each factor. The massive parallel processing allows us to complete all analyses within a day.

By MultiDCoX formulation, we identify DCX gene sets exhibiting B-type coexpression only [3]. The other two types of differential co-expression may be identified using multivariate differential expression analysis followed by clustering. MultiDCoX algorithm can be applied to different clinical data to quantify the influence of multiple cofactors on the co-expression and its associated phenotypes. Multiple aspects of the formulation and the algorithm need to be studied in our future improvements: Robustness of A_mn_(I) to outliers is an important aspects of the performance of the algorithm and impact of the thresholds used in the algorithm also to be studied. However, without tuning, the choice of parameters appears to be effective enough for both simulated and real data sets.

The application of MultiDCoX on a breast cancer data has revealed interesting sets of DCX genes: the set of *ESR1,* its cofactors along with downstream genes of ESR1 and genes associated with relevant ESR1 dependent transcriptional regulation; the set of genes containing ER binding site in their *cis* region. Furthermore, we have shown that the co-expression of gene sets that contain *CXCL13* and the gene sets that contain MMP1 is affected by ER status too in addition to tumor grade which couldn’t have been elicited in a typical univariate DCX analysis.

## Declarations

### Abbreviations

DCX: Differential Co-expression/Differentially Co-expressed
DE: Differential Expression
GO: Gene Ontology
KEGG: Kyoto Encyclopaedia of Genes and Genomes
FPs: False Positives
FNs: False Negatives
FNR: False Negative Rate
FDR: False Discovery Rate
FPR: False Positive Rate
OR: Odds Ratio
HPC: Hi Performance Computing
ER: Oestrogen Receptor

## Ethical Approval and Consent to participate

Not applicable

## Consent for publication

All authors read and approved the manuscript for submission.

## Availability of supporting data

The breast cancer dataset is available from the publication Miller *et al.* [25].

Analysis results are available as ***Additional File 1*:***UppsalaBCResults.xlsx*

## Acknowledgements

The authors would like to thank DrJoshy George, Dr. Foo Jia Nee and Dr. Astrid Irwanto for their helpful comments on the manuscript. We thank Swarna for helping in the early discussion of the project, DrsJuntao, Sigrid and Huaien for comments and discussion.

## Authors' contributions

RKMK conceived the project. RKMK and JCR guided HL to code the algorithm and carry out analysis. HL coded the algorithm. Both RKMK and HL drafted the manuscript and carried out all analyses. JCR revised the manuscript. All authors read and approved the final manuscript.

## Authors' information

HL: Genome Institute of Singapore, Singapore.

JCR: Nanyang Technological University, Singapore.

RKMK: The Jackson Laboratory, USA.

## Additional files

**Additional File 1: UppsalaBCResults.xlsx**

- Format: XLSX
- Title of Data: Results of Analysis of Breast Cancer Data
- Description: Contains all differentially co-expressed genesets with respective differential co-expression model fit (F-test p-value, coefficient value), gene counts, and permutation results over three factors (ER, p53 and Grade) in breast cancer data. Remarks: Grade+ indicates higher grade tumor i.e. 2 and 3, while Grade– indicates lower grade tumour i.e. 1.

